# Exploratory profiling of defense system signatures in *Klebsiella pneumoniae* clonal populations

**DOI:** 10.64898/2026.08.13.739500

**Authors:** Xiaofu Wan, Liyin Ji, Shuhong Han, Zixun Lin, Yukun Zeng, Min Ming, Wenheng Gan, Xiangke Duan, Hongzhou Lu, Jiayin Shen

## Abstract

Bacterial defense systems against bacteriophages are critical for bacterial genome stability and fitness, yet their distribution and epidemiological relevance in *Klebsiella pneumoniae* remain unexplored. We aimed to characterize defense system signatures across major *K. pneumoniae* clonal lineages and evaluate their utility as genomic signatures for tracking clonal dissemination. Here, we analyzed 6,346 genomes to characterize population structure, defense system signatures, and their coevolution with resistance determinants and plasmid backbones. PopPUNK clustering resolved 146 lineages that stratified into major clonal lineages (G-ST-KL combinations) with distinct resistance and virulence profiles. DefenseFinder and PADLOC identified 320 distinct defense systems, revealing that each clonal lineage harbors a unique defensotype characterized by systematic module replacement rather than stochastic gene loss. Co-occurrence networks further showed that these systems are organized into lineage-specific functional modules, with contrasting architectures even among lineages sharing the same sequence type. Integration of defense, resistance, and plasmid data uncovered strong lineage-specific associations, whereby broad-host-range plasmid backbones acquired distinct defense-resistance payloads in different clonal backgrounds. Geographic and host-niche analyses demonstrated that defense system distribution reflects clonal lineage expansion rather than independent geographic selection, and analysis of 689 Chinese genomes confirmed vertical inheritance of lineage-specific signatures along transmission chains. Collectively, defense systems in *K. pneumoniae* are organized into lineage-specific defensotypes shaped by synergistic modules and clonal evolutionary dynamics. Defense system profiling provides an additional layer of epidemiological resolution beyond conventional typing and offers practical utility for genomic surveillance, particularly in resource-limited settings where PCR-based detection of conserved systems could serve as a rapid proxy for identifying high-risk *K. pneumoniae* clones.

## Introduction

*Klebsiella pneumoniae* is a globally critical opportunistic pathogen and a leading cause of hospital-acquired infections, including pneumonia, urinary tract infections, bacteremia, and pyogenic liver abscesses. The emergence and pandemic spread of multidrug-resistant (MDR) and hypervirulent *K. pneumoniae* (hvKP) clones, particularly carbapenem-resistant *K. pneumoniae* (CRKP), have severely limited therapeutic options and posed an urgent threat to public health [1,2]. Central to its epidemic success is the remarkable capacity of *K. pneumoniae* to acquire and maintain mobile genetic elements, particularly resistance plasmids and integrative conjugative elements, that disseminate carbapenemases, extended-spectrum β-lactamases, and hypervirulence determinants across clinical, environmental, and animal reservoirs [3,4]. Yet, the evolutionary forces that govern the stability and transmission of these elements within *K. pneumoniae* populations remain incompletely understood. In China, ST11 of *K. pneumoniae* has emerged as the predominant CRKP clone, with capsular types KL64 and KL47 being the most prevalent. Notably, ST11-KL64 strains have gradually displaced ST11-KL47 strains as the dominant clone, with detection rates increasing from 28.2% in 2016 to 45.7% in 2020 [5,6,7]. These high-risk clones frequently acquire hypervirulence plasmids, evolving into carbapenem-resistant hypervirulent *K. pneumoniae* (CR-hvKP) “dual-threat” pathogens that combine extensive antimicrobial resistance with enhanced virulence [8]. ST15 is an emerging high-risk clone in China, with pan-drug-resistant strains reported, displaying a complex carbapenemase spectrum alongside high ESBL and fluoroquinolone resistance [9,10]. ST23 is the classic hypervirulent lineage causing fatal community-acquired infections; it has recently acquired carbapenemase genes, leading to CR-hvKP emergence, and possesses the most extensive virulence arsenal. ST258/ST412 also carries multiple carbapenemases, confirming their global dissemination as high-risk clones [11–15].

One critical but underappreciated force shaping bacterial genome evolution is phage predation. The ability of lytic bacteriophages to rapidly devastate bacterial populations has driven the evolution of multiple layers of defense, including a diverse array of specialized anti-phage defense systems [16]. In *Vibrio cholerae*, phage predation has been shown to limit epidemic duration and severity, while bacteria acquire anti-phage defense systems through mobile elements such as SXT/R391 integrative conjugative elements that simultaneously carry antibiotic resistance genes, creating mobile defense-resistance cassettes wherein antibiotic resistance and phage defense genes are physically linked and co-transmitted [17,18]. This co-evolutionary dynamic exemplifies the intimate connection between phage defense and the ecology of mobile genetic elements [19]. For *K. pneumoniae*, the interplay between phage pressure and the stability of resistance plasmids represents a significant knowledge gap. This is particularly pertinent given the global dissemination of high-risk clones such as ST11 (carbapenem-resistant), ST15 (ESBL-producing), and ST23 (hypervirulent), often associated with the acquisition of specific plasmid types [20,21]. It remains unknown whether these clones carry lineage-specific defense profiles.

Over 150 distinct anti-phage defense systems have been identified across the microbial pangenome, revealing extraordinary mechanistic diversity including abortive infection, nucleotide depletion, membrane protection, and direct phage DNA degradation [22]. Wu and Garushyants et al. reveal that defense systems in *E. coli* strains are generally compatible and, in some instances, interact resulting in synergistic antiphage effects, conferring an evolutionary advantage on bacteria under specific environmental conditions [23]. In contrast, our laboratory recently investigated clinical *P. aeruginosa* isolates and found that the number of defense systems per strain does not correlate with its intrinsic phage resistance. Instead, we identified specific pairs of defense systems that exhibit additive, synergistic, or neutral interaction outcomes, highlighting that functional relationships between systems are a critical determinant of the anti-phage defense landscape [24]. While a few small-scale studies have examined individual defense systems in *K. pneumoniae* lineages--for instance, Kadkhoda et al. linked CRISPR-Cas carriage to *bla*NDM-1, biofilm formation, and virulence genes in CRKP [25]. However, these investigations have been geographically and numerically limited, leaving the population-level defense system landscape across epidemic clones entirely unexplored.

Here, we systematically profiled defense system repertoires across 6,346 globally distributed *K. pneumoniae* genomes to investigate whether phage defense architectures are structured by clonal background, vertically inherited, and coevolved with resistance plasmids. By integrating population genomics with defense system annotation, resistance profiling, and plasmid replicon typing, we assessed the lineage specificity and modular organization of defense signatures and their association with the resistance mobilome. We further evaluated the geographic and host-niche distribution of these systems and leveraged transmission network analysis to test their utility as molecular markers for tracking clonal dissemination. This work establishes a framework for understanding the interplay between phage defense, antimicrobial resistance, and plasmid evolution, with implications for genomic surveillance of high-risk *K. pneumoniae* lineages.

## Result

### Population structure and clonal architecture of *K. pneumoniae*

To elucidate population structure, we performed PopPUNK clustering on 6,346 *K. pneumoniae* genomes from PATRIC (1982-2024, 60 countries) (**Table S1**). PopPUNK resolved 146 lineages, with Cluster 1 comprising 71% (4,512/6,346) of isolates (**Figure S1**) (**Table S2**). Sub-clustering stratified Cluster 1 into five groups: G1A (n=1,337), G1B (n=1,068), G1C (n=821), G1D (n=652), and G1E (n=634). The remaining isolates were consolidated into G2 (Cluster 2-25, n=1,049) and G3 (Cluster 26-146, n=785) (**Figure 1A)**. Molecular typing revealed marked ST and KL differentiation across these groups **(Figure 1B**) (**Table S3**). G1A and G1B were overwhelmingly dominated by ST11 (93.6% and 94.4%), but exhibited divergent capsule profiles: KL64 was enriched in G1A (83.4%), whereas KL47 predominated in G1B (59.4%). G1C was primarily ST15 (70.3%), associated with KL112 and KL19. G1D was dominated by ST37 and ST17. G1E comprised mainly ST23 (41.3%) and ST147 (20.4%), with KL1 (43.7%) as the dominant capsule type. G2 was more diverse, harboring ST258/ST412 and KL2/KL57. G3 contained various rare STs and a broad range of KL types. Correspondingly, these groups showed significant differences in resistance and virulence gene loads (**Figure 1C&D**), consistent with their distinct clinical threat profiles. G1A/G1B (predominantly ST11) dominate carbapenem-resistant *K. pneumoniae* (CRKP), carrying KPC-2 in >87% of isolates and yersiniabactin in >90%; G1A additionally harbors aerobactin and *rmpA*, reflecting ongoing evolution toward CR-hvKP dual-threat clonal lineages (**Figure S2**). G1C (mainly ST15) represents an emerging high-risk group with drug-resistant strains, displaying complex carbapenemase profiles (KPC-2, OXA-232) alongside high ESBL and fluoroquinolone carriage. G1E (dominated by ST23 and ST147) represents the classical hypervirulent lineage causing fatal community-acquired infections; recent acquisition of carbapenemase genes has led to CR-hvKP emergence, and it possesses the broadest virulence arsenal (aerobactin, yersiniabactin, colibactin, salmochelin, *rmpA*). G2 (harboring ST258/ST412) carries diverse carbapenemases (KPC, NDM, OXA-48-like), confirming their global spread as high-risk strains. G1D and G3 showed lower overall resistance and virulence, serving as comparative controls.

**Figure 1.**
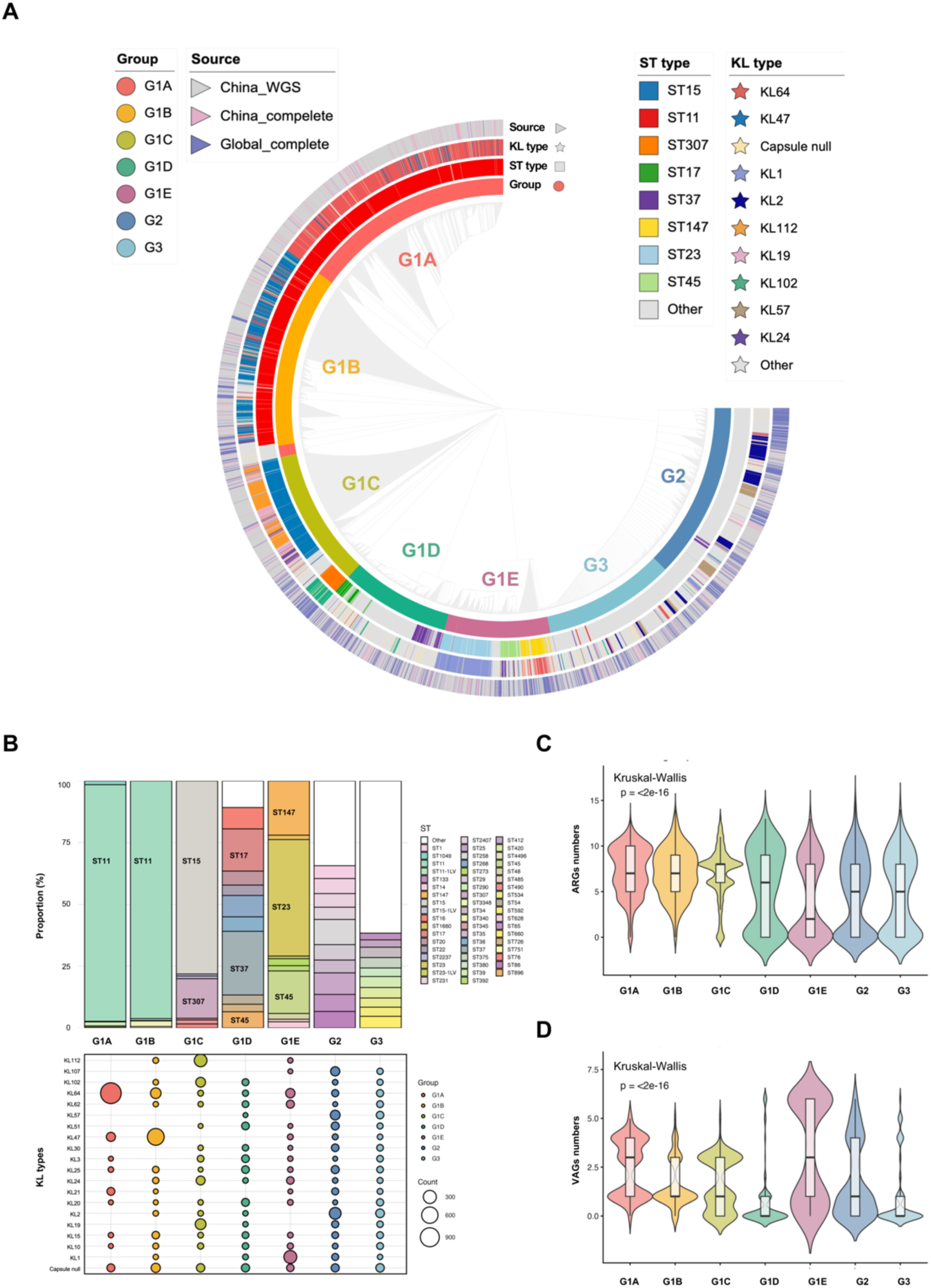
Population structure and clonal architecture of *K. pneumoniae*. **(A)** Hierarchical clustering tree of 6,346 *K. pneumoniae* genomes based on PopPUNK analysis. The tree was rooted at the midpoint and annotated with external rings indicating group assignment (G1A-G3), isolation source, ST type, and KL type. **(B)** Stacked bar plots showing the distribution of sequence types (STs) within each group (G1A-G3). Bubble plot depicting the prevalence of capsular types (KLs) within each group, with bubble size proportional to the prevalence rate. **(C)** The distribution of resistance gene counts across the seven groups. Statistical significance was assessed using the Kruskal-Wallis test (p < 2e -16 for comparisons). **(D)** The distribution of virulence gene counts across the seven groups. Statistical significance was assessed using the Kruskal-Wallis test (p < 2e -16 for comparisons).

Capsule distribution within each group-ST combination revealed strict ST-KL lock-in patterns (**Table S4**): ST23 with KL1, ST11 with KL64 in G1A but KL47 in G1B, ST307 with KL102, and ST15 with KL112. Therefore, we defined clonal lineages as unique G-ST-KL combinations, based on the strict ST-KL lock-in patterns observed across the population. For subsequent major analyses, we identified G1A-ST11-KL64 (n=1,107), G1B-ST11-KL47 (n=628), G1C-ST15-KL112 (n=274), G1C-ST307-KL102 (n=103), G1C-ST15-KL19 (n=149), G1E-ST23-KL1 (n=255), and G1E-ST147-KL64 (n=75) as the primary clonal lineages of *K. pneumoniae* based on their high prevalence and distinct clinical threat profiles.

### Defense system distribution of *K. pneumoniae* is lineage-specific

To characterize defense system variation across clonal lineages, we identified 320 distinct defense systems using DefenseFinder and PADLOC across 6,346 *K. pneumoniae* genomes (**Figure 2**) (**Table S5)**. Strains carried 17-30 defense systems, with G1A, G1B, and G1C harbored the highest numbers, whereas G1E, G2, and G3 carried fewer (Kruskal-Wallis, *p* < 0.001; **Figure S3**). We next defined defense system signatures for the major clonal lineages (**Table S6**). Strikingly, the defense system signatures were not merely quantitatively distinct but exhibited systematic module replacement, with each clonal lineage displaying a unique combination of core and accessory systems (**Figure 3A**). G1A-ST11-KL64 and G1B-sST11-KL47 shared a conserved defense core (PD.T4.3, AbiE, PDC.S13, Mok_Hok_Sok, RM_Type_IV), with G1A additionally enriched for PDC.S15, reflecting fine-tuning within the ST11 backbone across different group backgrounds. G1C-ST15-KL112/KL19 displayed a complementary signature (RosmerTA, Zorya_TypeI, UG4, Hachiman, CBASS_I) while entirely lacking the G1A/B core, indicating wholesale replacement of defense architecture. G1C-ST307-KL102 exhibited ST307-specific enrichment (Hypnos, Rst_TIR.NLR, Shango) and complete absence of ST15-characteristic systems, demonstrating that ST-KL swaps within the same group yield distinct defense system signatures. G1E-ST23-KL1 possessed a unique signature (DISARM_1, DRT_2, Mokosh_Type_I_A) but entirely lacked AbiE, a system universally present in G1A-G1D, whereas G1E-ST147-KL64 retained AbiE and RM_Type_IV while acquiring DS.32 and Druantia_III, yielding a hybrid signature closer to G1A/B. These lineage-specific absence patterns suggest systematic module replacement rather than stochastic gene loss, as exemplified by the RosmerTA-Zorya-Hachiman-UG4 module in G1C-ST15-KL112/KL19, the Hypnos-Rst-Shango module in G1C-ST307-KL102, and the DISARM-DRT-Mokosh module in G1E-ST23-KL1.

**Figure 2.**
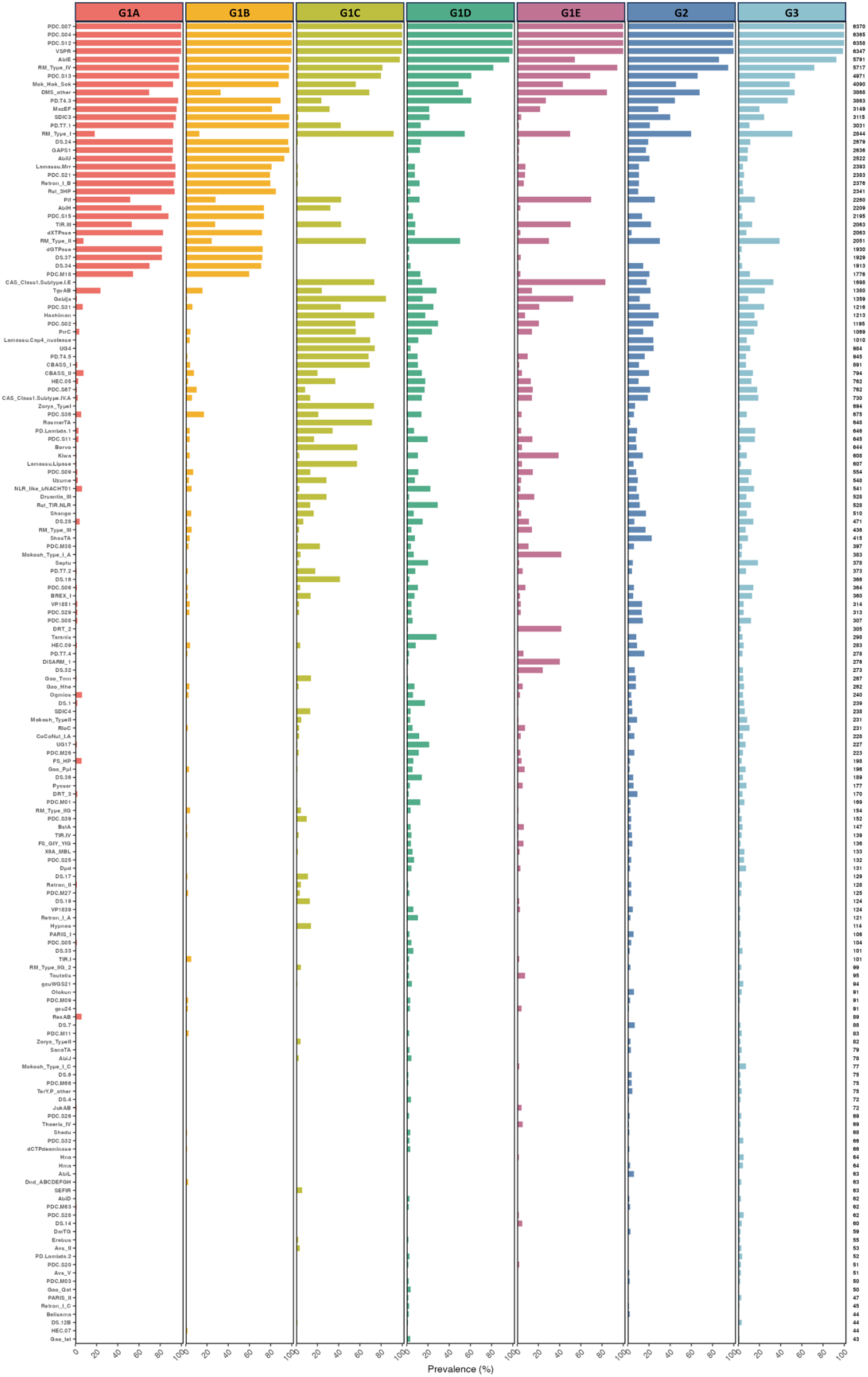
Prevalence of defense systems across seven groups. Bar plot showing the prevalence (percentage of isolates carrying each system) of 320 distinct defense systems identified by DefenseFinder and PADLOC across the seven groups (G1A-G3).

**Figure 3.**
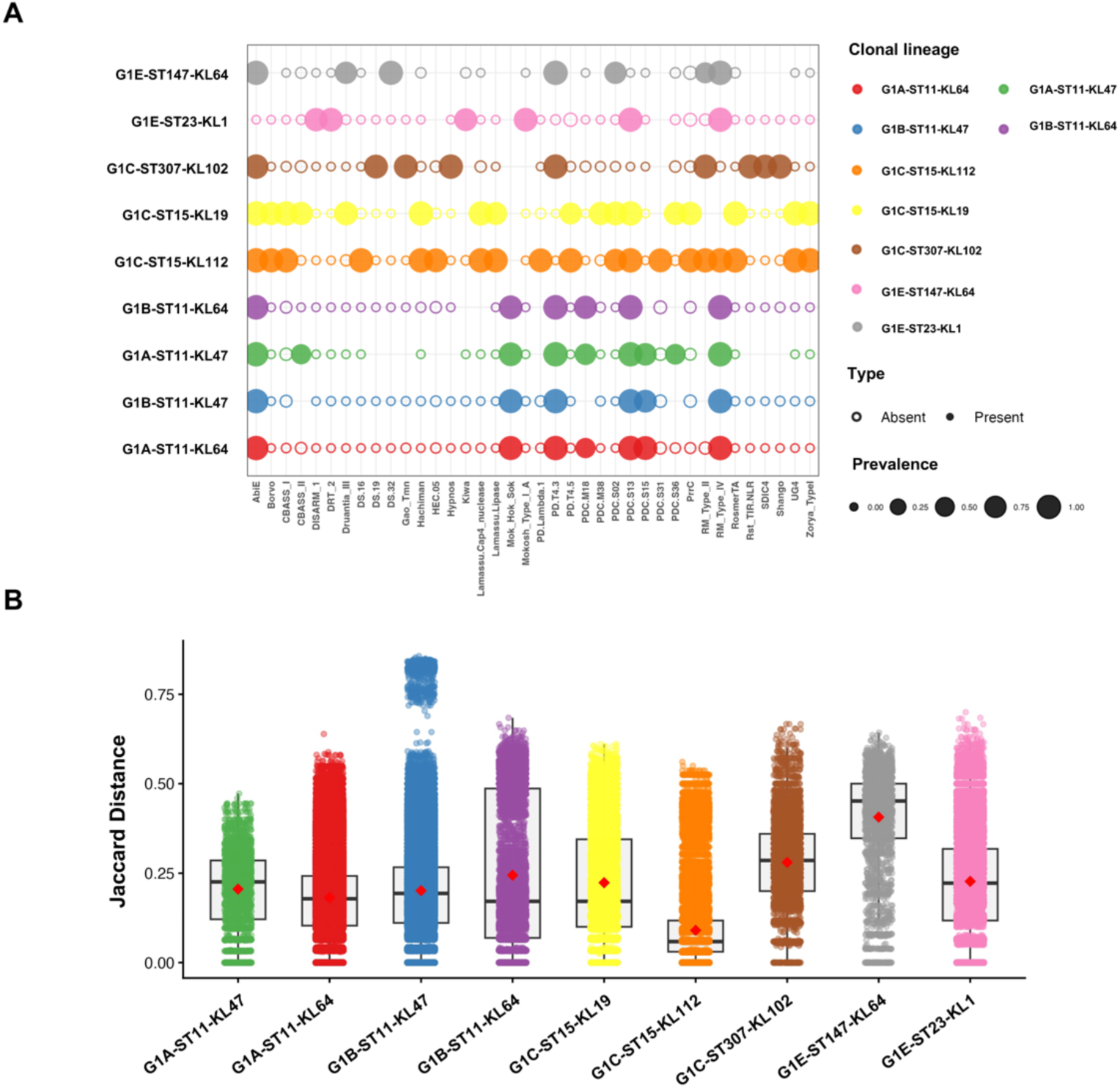
Lineage-specific defense system signatures and compositional conservation. **(A)** Bubble plot showing the prevalence of 38 selected defense systems across the nine major clonal lineages. Bubble size is proportional to the prevalence rate within each lineage. White circles indicate absence of a given system, while colored circles indicate presence, with color intensity reflecting the prevalence level. **(B)** Box plot showing the distribution of pairwise Jaccard distances within each of the nine major clonal lineages.

To evaluate the compositional conservation of defense systems across different taxonomic levels, we calculated pairwise Jaccard distances **(Table S7**) among strains within each group and further assessed internal homogeneity using the coefficient of variation (CV = standard deviation / mean) (**Table S8**). Overall, the conservation of defense systems increased markedly with finer taxonomic resolution. At the group level (**Figure S4A**), Jaccard distances ranged from 0.248 (G1A) to 0.727 (G3), with a median of approximately 0.54. At the ST level (**Figure S4B**), the median distance decreased to 0.34 (from 0.239 for ST23 to 0.622 for ST37). At the clonal lineage level (**Figure 3B**), the median distance further declined to 0.22, with the majority of lineages (8/9) exhibiting average distances below 0.30; among these, G1A-ST11-KL64 (0.182) and G1B-ST11-KL47 (0.201) showed the highest compositional consistency. This hierarchical trend indicates that defense system signatures are most conserved at the clonal lineage level, supporting their potential as markers for lineage identification. However, considerable variation in within-lineage homogeneity was observed, reflecting differences in evolutionary dynamics among lineages. CV values at the clonal level ranged from 0.36 (G1E-ST147-KL64) to 1.18 (G1C-ST15-KL112). Notably, G1C-ST15-KL112, despite having an extremely low average distance (0.090), exhibited a CV as high as 1.18, indicating substantial dispersion in its internal distance distribution (**Figure S4C**). The distribution of pairwise distances (**Figure S4D**) was markedly right-skewed, with the vast majority of values clustering below 0.2 and a few extending to 0.4 or above. This suggests that the defense system signatures of most strains within this clonal lineage are nearly identical, but a few strains deviate considerably, possibly due to horizontal gene transfer or mixed typing. Joint analysis of average distance and CV enabled effective identification of robust markers; for instance, G1A-ST11-KL64 and G1B-ST11-KL47, which combine high conservation with low internal variability, represent preferred candidates for clonal lineage identification.

### Functional assembly and co-occurrence networks of defense systems

Co-occurrence network analysis confirmed that defense systems are organized into functional modules (**Table S9 & Table S10**). G1A-ST11-KL64 (**Figure 4A**) is primarily supported by two stable high-frequency modules: a membrane barrier/innate immune signaling module (DS.34, DMS_other, TIR.III, Pif, PDC.M18) and a dNTP depletion module (dGTPase, DS.37, dXTPase, AbiH). G1B-ST11-KL47 (**Figure 4B**), despite sharing the ST11 backbone with G1A, exhibits a fundamental resetting of its defense strategy: it retains the dNTP depletion core but replaces the membrane barrier module with PD.T4.3-Lamassu.Mrr. G1C-ST15-KL19 (**Figure 4C**) shifts toward a CBASS-Gabija-VSPR module, whereas G1C-ST15-KL112 (**Figure 4D**) deploys a 12-system module supplemented by an independent Uzume module, representing the largest defense arsenal among all lineages. This contrast demonstrates that different KL types within the same ST can evolve entirely distinct module architectures. G1E-ST23-KL1 (**Figure 4E**) retains only two streamlined modules: a DRT_2-Mokosh_Type_I_A-Gabija-DISARM_1 core module and a PDC.S31-Mok_Hok_Sok-MazEF toxin-antitoxin module. Moreover, G1C-ST307-KL102 was centered on Retron and TIR-NLR, and G1E-ST147-KL64 exhibited a chimeric architecture combining G1A-like, G1C-ST15-like, and G1C-ST307-like systems (**Table S11**).

**Figure 4.**
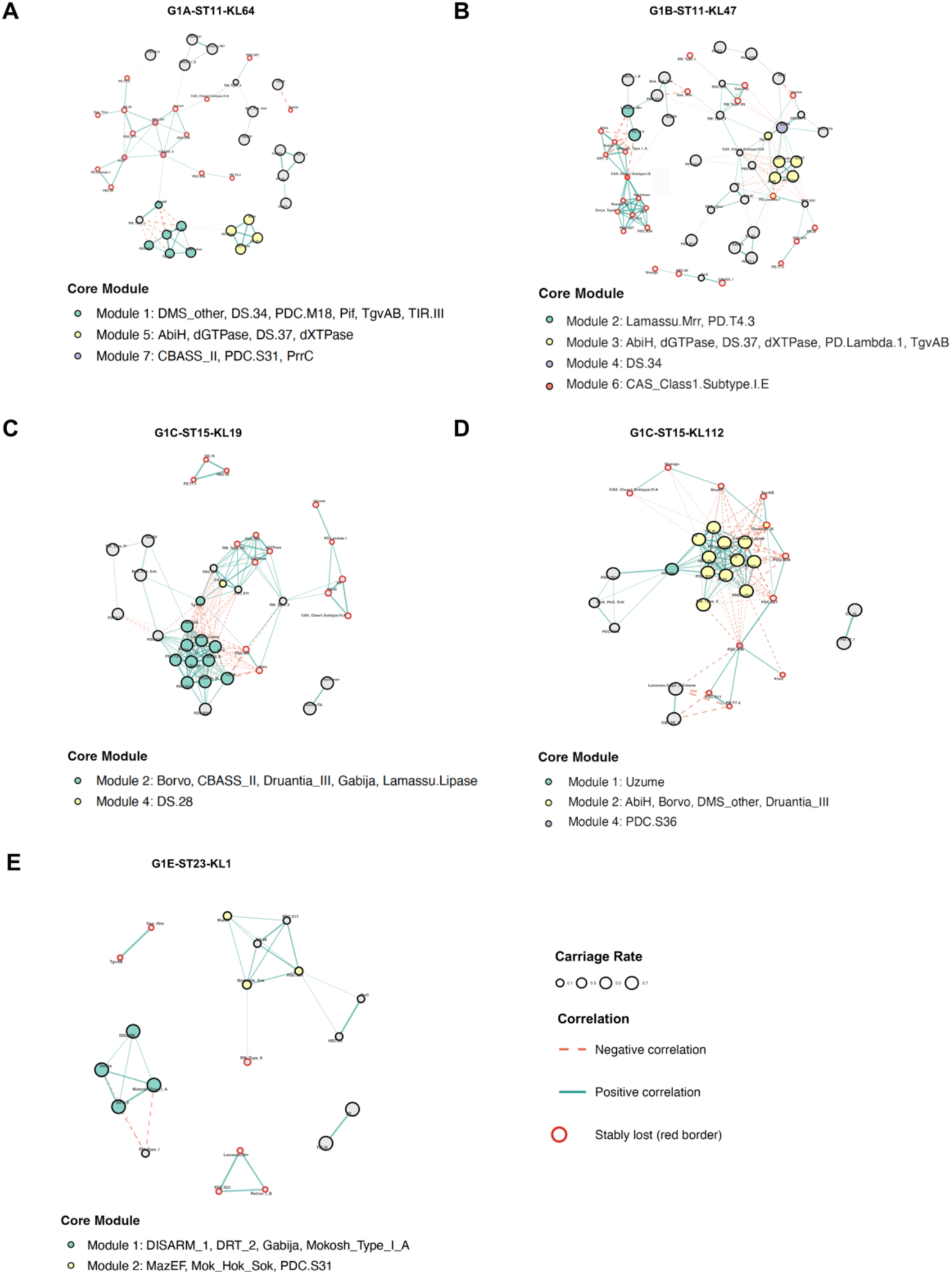
Co-occurrence networks of defense systems across major clonal lineages. **(A-G)** Co-occurrence networks showing significant pairwise correlations (|Phi| ≥ 0.2, Q < 0.05) among defense systems within each of the seven major clonal lineages: G1A-ST11-KL64 (A), G1B-ST11-KL47 (B), G1C-ST15-KL19 (C), G1C-ST15-KL112 (D), G1C-ST307-KL102 (E), G1E-ST23-KL1 (F), and G1E-ST147-KL64 (G). Nodes represent individual defense systems, with node size proportional to the system carriage rate within the clonal lineage. Node colors indicate core module assignments, with gray nodes representing systems not assigned to any core module. Red node borders denote systems with stable loss (prevalence < 5%). Edges represent significant pairwise correlations: solid cyan edges indicate positive co-occurrence, while dashed orange edges indicate negative co-occurrence (mutual exclusion). Edge thickness is proportional to the absolute Phi coefficient.

Across all lineages, stable loss patterns further corroborated the modular framework. In G1A-ST11-KL64, CBASS_II together with its downstream effectors PDC.S31, PrrC, Uzume, and PDC.S11 formed a complete low-frequency residual module (Module7), indicating that the canonical CBASS cascade has been systematically lost in the vast majority of isolates. In G1B-ST11-KL47, CAS_Class1.Subtype.I.E together with UG4, Hachiman, PD.T4.5, RosmerTA, and Zorya_TypeI constituted another low-frequency co-occurrence module (Module6). G1C-ST15-KL19 lost TIR.III and Pif, whereas G1C-ST15-KL112 lost only MazEF, revealing contrasting loss patterns even within the same ST15 background. Additionally, AbiE was exclusively absent in G1E-ST23-KL1. These lineage-specific absence patterns, rather than stochastic deletions, further reinforce the systematic nature of module replacement and suggest that defense system composition holds potential as a molecular marker for clonal lineage identification.

### Associations of defense systems with resistance genes and plasmid types

The lineage-specific assembly patterns of defense modules suggest functional implications beyond genome architecture. To explore this, we integrated defense modules, resistance genes, and plasmid dissemination into a unified framework (**Table S12 & Table S13**). At the strain level, defense-resistance co-occurrence exhibited strong lineage specificity (**Figure 5A**). In G1A-ST11-KL64, the membrane barrier module (DS.34, DMS_other, TIR.III, Pif, PDC.M18) positively correlated with sulfonamide (sul2), quinolone (qnrS1), and tetracycline (tet(A)) resistance (Phi = 0.28-0.51), but negatively correlated with aminoglycoside (aph3-Ia), macrolide (mphA, Mrx), and trimethoprim (dfrA12) resistance (Phi = -0.40 to -0.64). In G1B-ST11-KL47, CBASS_I showed highly significant co-occurrence with ermB (Phi = 0.846), while TgvAB and CBASS_II associated with dfrA1, qnrS1, and tet(A). The dNTP depletion module (dGTPase, DS.37, dXTPase, AbiH) consistently negatively correlated with catB3 across both ST11 backgrounds. In G1C-ST15-KL112, the super-module members (AbiH, PrrC, PDC.S31, Lamassu.Lipase, Borvo, etc.) positively associated with OXA-232 and rmtF but negatively with KPC-2 and sul1, revealing intra-clonal subpopulation differentiation. Notably, the G1C group-specific signature systems (RosmerTA, Zorya_TypeI, UG4, Hachiman) showed no resistance associations. In G1E-ST23-KL1, Mok_Hok_Sok, PDC.S31, and PDC.S11 showed exceptionally strong correlations with arr-3, dfrA27, aadA16, and KPC-2 (Phi ≥ 0.95), suggesting strong lineage-associated co-occurrence, potentially reflecting chromosomal linkage or clonal background (**Table S14**).

**Figure 5.**
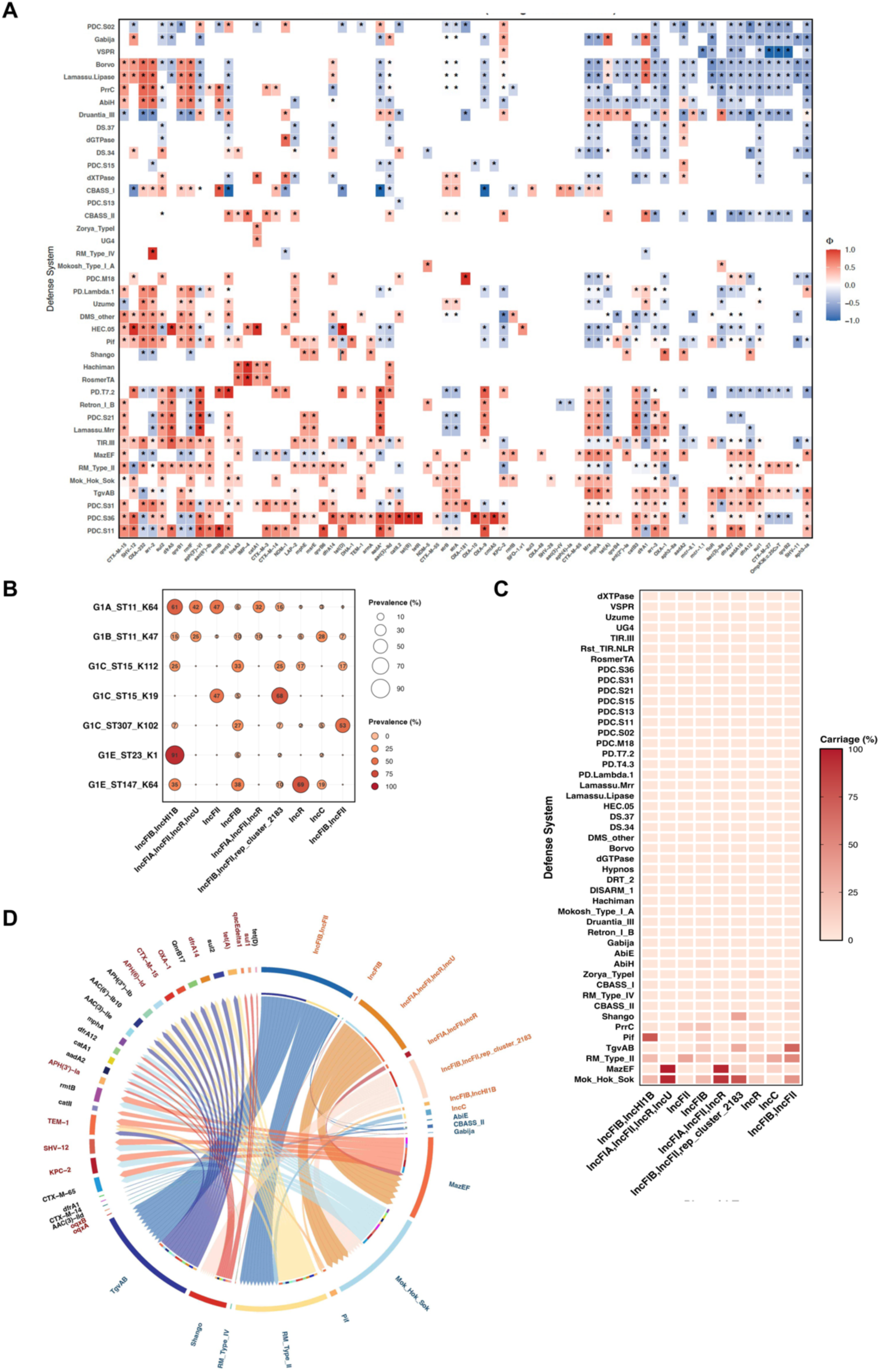
Associations of defense modules with resistance genes and plasmid dissemination. **(A)** Co-occurrence heatmap showing significant pairwise associations (Phi coefficient) between defense modules/systems and resistance genes across all the *K. pneumoniae* strains. Only associations with |Phi| ≥ 0.2 and Q < 0.05 are shown. Cells are color-coded from blue (negative association) to red (positive association), with color intensity proportional to Phi. *Asterisks indicate the significance level after FDR correction: *Q < 0.05. **(B)** Bar plot showing the prevalence of major plasmid clusters across the seven clonal lineages. **(C)** Carriage rate analysis showing the prevalence of defense systems on plasmid clusters. **(D)** Circos plot illustrating the integrated plasmid-defense-resistance network. The three concentric sectors represent plasmid clusters (orange), defense systems (blue), and resistance genes (red), respectively. Links between sectors indicate physical co-carriage on the same plasmid backbone, with link width proportional to the co-carriage frequency.

At the plasmid level, plasmid prevalence varied markedly across lineages, with IncFIB/IncHI1B plasmid cluster strikingly enriched in G1E-ST23-KL1 clonal lineage (90.6%). The G1E-ST147-KL64 clonal lineage displayed the most diverse plasmid repertoire (IncR 69.2%, IncFIB 38.5%, IncFIB/IncHI1B 34.6%) (**Figure 5B)**. Carriage rate analysis on plasmid clusters showed that most defense systems were nearly absent from plasmids, reinforcing their chromosomal localization (**Figure 5C)**. The G1C-specific signature systems (RosmerTA, Zorya_TypeI, UG4, Hachiman) were almost entirely absent from plasmids, with exceptions including Shango (a G1C-ST307-KL102 specific system) on IncFIB/IncFII/rep_cluster_2183 (36.1%) and AbiE and Retron_I_B on rep_cluster_1254 (75%). Only a subset of strain-level associations was recapitulated on plasmids: CBASS_II-dfrA1 remained significant, confirming plasmid mediation (**Figure S6**). In contrast, CBASS_II associations with qnrS1 and tet(A) were not significant at the plasmid level (|Phi| < 0.2), indicating chromosomal co-inheritance for these partners. Similarly, the CBASS_I-ermB association showed no plasmid signal, supporting its chromosomal origin. The plasmid-defense-resistance network centered on the IncFIB/IncFII/rep_cluster_2183 plasmid cluster, which carried different defense and resistance payloads in different lineages (**Figure 5D & Table S15**): in G1C-ST307-KL102, it linked Shango with OXA-1, aadA2, armA, and msrE; in G1B-ST11-KL47, the same plasmid type linked TgvAB and CBASS_II with dfrA1. This indicates that IncFIB/IncFII/rep_cluster_2183 acts as a broad-host-range plasmid backbone capable of acquiring different defense-resistance payloads in different clonal backgrounds. Network analysis revealed that the IncFIB/IncHI1B plasmid cluster linked Mok_Hok_Sok with sul2, qnrS1, and tet(A), a set distinct from its chromosomal partners (arr-3, dfrA27, aadA16, KPC-2), suggesting that Mok_Hok_Sok cooperates with different resistance genes in different genetic compartments. This association was supported by carriage rate data: Mok_Hok_Sok was carried on IncFIB/IncHI1B at 90.6% in G1E-ST23-KL1, making it the primary plasmid vector for Mok_Hok_Sok in this clonal lineage. Additionally, Mok_Hok_Sok showed substantial carriage on IncFIB (24.3%) and IncFIB/ IncFII/rep_cluster_2183 (44.4%), indicating its association with multiple plasmid backbones.

### Global distribution patterns and host niche characteristics of defense system signatures

To assess whether defense systems exhibit geographic preferences, we analyzed the geographic distribution of strains (**Table S16**). The results showed that AbiE, as a species-core system, maintained stable prevalence rates of 85%-95% across all continents without any significant geographic clustering, reinforcing its role as a conserved backbone component of the *K. pneumoniae* pan-genome. In contrast, G1C-lineage characteristic systems (RosmerTA, Zorya_TypeI, Hachiman, UG4, etc.) and G1E-ST23-KL1 characteristic systems (DISARM_1, DRT_2) were significantly enriched in the Asia-Pacific region, while Europe and North America exhibited delayed enrichment and fluctuating distributions (**Figure 6A & Figure 6B**). The prevalence of defense systems also varied significantly across different isolation sources (**Figure 6C**). AbiE maintained extremely high and stable prevalence across all hosts (82.4%-100%). PDC.S15 was exceptionally enriched in environmental isolates (78.3%), far exceeding that in human (39.5%), other (15.2%), and animal (5.0%) sources, further supporting the environmental reservoir potential of the G1A-ST11-KL64 clonal lineage; together with its dominance in the transmission network (82.4% of links), this suggests continuous dissemination through the environment-human interface. DISARM_1 and DRT_2 showed higher prevalence in “other” sources (11.3% and 11.1%, respectively) than in human sources (3.8% and 4.4%), implying the existence of non-human reservoirs. Meanwhile, CAS_Class1.Subtype.I.E was more prevalent in animal (29.4%) and other (31.4%) sources than in human (25.8%); Hachiman was mainly concentrated in human and environmental sources (19.9% and 16.8%) but completely absent in animals. Collectively, defense system distribution is shaped both by clonal background and by fine-tuned niche selection; enrichment of certain systems in environmental or non-human sources suggests potential reservoirs, whereas AbiE serves as a universal backbone that remains stable across diverse hosts.

**Figure 6.**
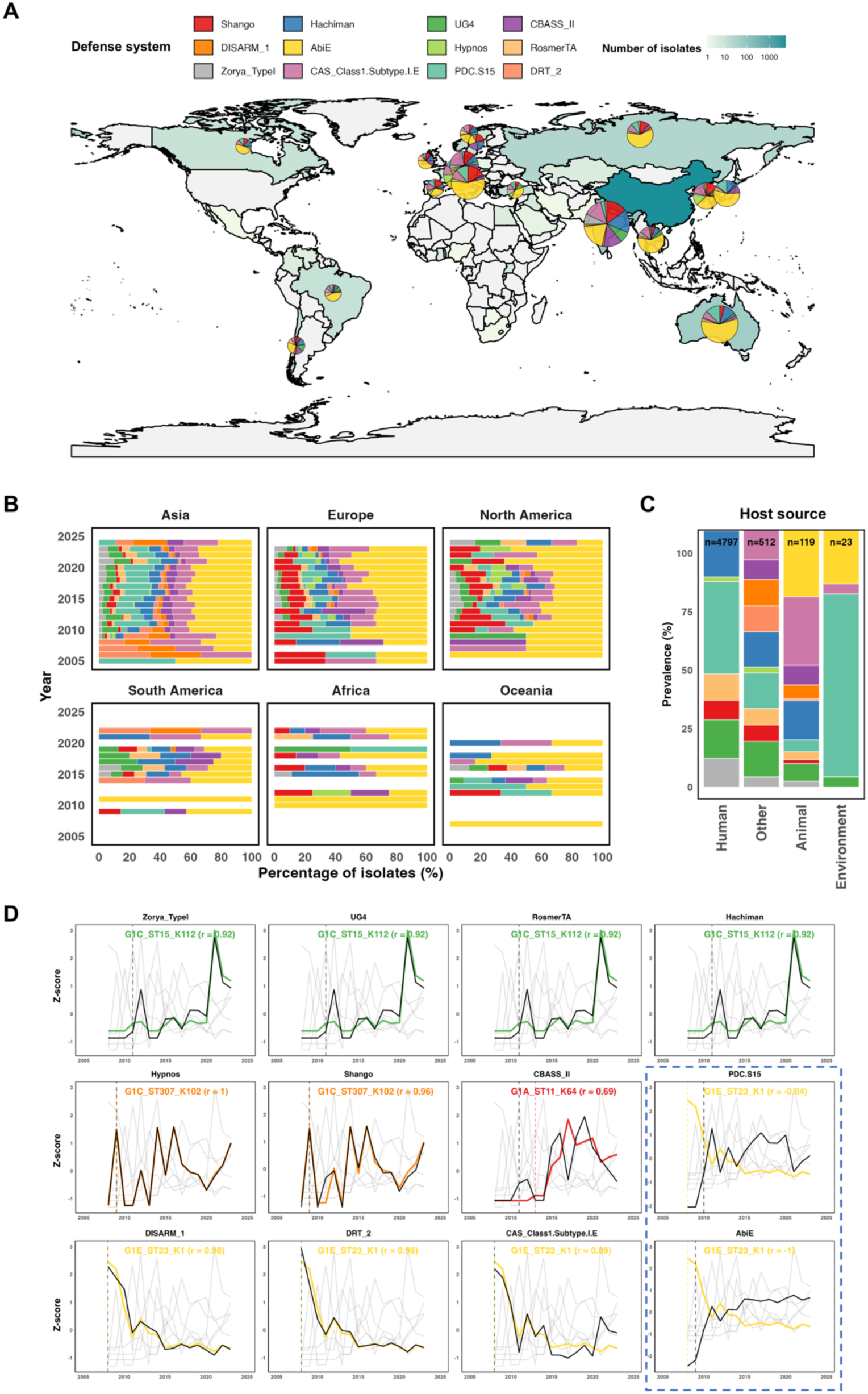
Global distribution, temporal dynamics, and host niche characteristics of defense system signatures. **(A)** World map showing the geographic distribution of defense systems across different countries. Pie charts overlaid on each country represent the prevalence (carriage rate) of key defense systems within that country, with chart size proportional to the number of isolates analyzed. Due to the exceptionally large number of isolates from China, pie charts are not displayed for China to avoid visual overcrowding. **(B)** Stacked area plots showing the temporal dynamics of defense system prevalence across six continents from 2005 to 2025. Each colored area represents the prevalence of a specific defense system or system group over time. **(C)** The prevalence of key defense systems across different isolation sources: human, animal, environmental, and other sources. **(D)** Time-series correlation plots showing the synchrony between defense system prevalence (black lines) and the contemporaneous proportions of their corresponding clonal lineages (colored lines, representing the best-matching clonal lineage). Both variables are standardized as Z-scores for direct visual comparison. Dashed vertical lines indicate the first detection year of the defense system (black) and its best-matching clonal lineage (color-matched). The best-matching clonal lineage and its Pearson correlation coefficient (r).

The geographic and host-level distribution differences described above suggested intrinsic associations between defense systems and specific clonal lineages. To directly test this hypothesis, we performed time-series correlation analysis between defense system prevalence and the contemporaneous proportions of each clonal lineage (**Figure 6D & Table S17**). The results confirmed a high degree of synchrony between defense system signatures and specific clonal lineages. Hachiman-UG4-RosmerTA-Zorya_TypeI showed strong positive correlations with the proportion of the G1C-ST15-KL112 clonal lineage (r = 0.927-0.939); Hypnos was almost perfectly synchronized with G1C-ST307-KL102 (r = 0.991); and DISARM_1/DRT_2 correlated with G1E-ST23-KL1 with coefficients of 0.976-0.997. Conversely, AbiE exhibited a strong negative correlation with G1E-ST23-KL1 (r = -0.994), indicating a systematic loss of this core system in the G1E lineage. Notably, although Shango was highly synchronized with G1C-ST307-KL102 (r = 0.935), its global prevalence (8.02%) was much higher than that of Hypnos (1.80%), suggesting that this defense system may possess stronger intercontinental transmission capacity, potentially via the ST307 clonal lineage or associated mobile genetic elements (e.g., the IncFIB/IncFII/rep_cluster_2183 plasmid, which links Shango with multiple resistance genes in ST307). Together, these results indicate that the spatiotemporal distribution of defense system signatures likely represents a direct reflection of clonal lineage geographic expansion.

### Vertical inheritance validation of defense system signatures along transmission chains within China

To further validate the vertical inheritance conservation of defense system signatures at a finer scale, we constructed a high-resolution transmission network based on 689 complete genomes from China (**Figure 7**). Analysis of inter-provincial transmission links (SNP ≤ 20) revealed that G1A-ST11-KL64 was the dominant circulating clonal lineage (**Table S18**). The strongest transmission corridors, Hubei-Sichuan (149 links), Hunan-Zhejiang (107 links), and Jiangxi-Zhejiang (62 links), all involved G1A-ST11-KL64 on both endpoints, indicating clonal expansion rather than sporadic transmission. Notably, geographically proximate provinces did not necessarily exhibit strong transmission links unless they shared the same clonal background: while Zhejiang-Shanghai showed 62 links, Zhejiang-Jiangsu showed only 15 links, and Zhejiang-Anhui showed only 7 links. This pattern demonstrates that transmission intensity is determined by clonal lineage rather than geographic distance alone. In Hubei, where G1A-ST11-KL64 clonal lineage comprised 97.1% of isolates, PDC.S15 carriage reached 96.88% and remained stable over time. Similarly, in Jiangxi and Zhejiang, PDC.S15 consistently paralleled the prevalence of G1A-ST11-KL64 clonal lineage. In contrast, in provinces with higher clonal diversity, such as Beijing and Shanghai, PDC.S15 was exclusively confined to ST11 clonal lineages (G1A/G1B), while non-ST11 clonal lineages (e.g., G1E-ST23-KL1) completely lacked PDC.S15. Notably, G1E-ST23-KL1 clonal lineage displayed an alternative defense architecture characterized by DISARM_1 and DRT_2, while AbiE was consistently lost in G1E-ST23-KL1 across all provinces. These patterns demonstrate that defense systems are stable clonal lineage-specific signatures that are faithfully transmitted along with specific clonal lineages, rather than being acquired through geographic dissemination. To quantitatively assess the vertical inheritance of these lineage-specific defense systems, we performed Pagel’s λ phylogenetic signal tests within each clonal lineage (**Table S19**). PDC.S15 exhibited exceptionally high λ values in G1A-ST11-KL64 (λ = 0.9999, P < 0.001) and G1B-ST11-KL47 (λ = 0.9999, P = 0.013), indicating strong phylogenetic conservation and vertical transmission within G1A-ST11-KL64 and G1B-ST11-KL47 clonal lineages. In contrast, Shango and DISARM_1 showed λ values near zero, consistent with horizontal transfer via mobile genetic elements. Collectively, these results establish defense system signatures as robust molecular markers for tracking *K. pneumoniae* clonal lineages.

**Figure 7.**
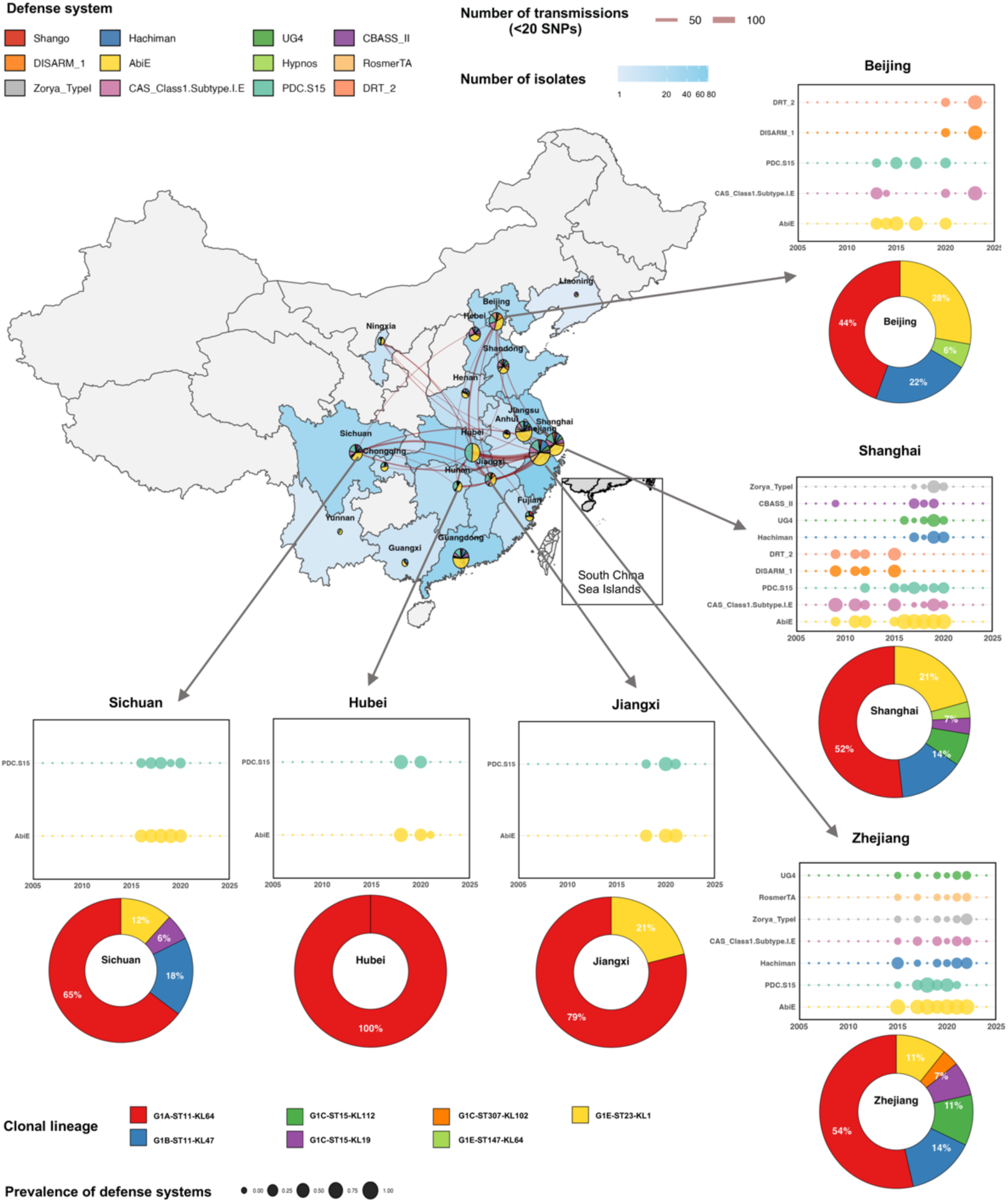
Geographic distribution, transmission network, and clonal lineage-specific defense system signatures of *K. pneumoniae* in China. The main map shows China with inter-provincial transmission networks among 689 *K. pneumoniae* complete genomes. Pie charts overlaid on each province represent the prevalence of 12 defense systems, with sector size proportional to the carriage rate. Red curves indicate inter-provincial transmission links (SNP ≤ 20), with line width proportional to transmission intensity (pair count). Arrows point to six key provinces (Beijing, Hubei, Jiangxi, Shanghai, Sichuan, and Zhejiang). For each of the six key provinces, two panels are displayed to the right or below the map. Bubble plots show the temporal dynamics of representative defense systems from 2005 to 2024, where bubble size is proportional to carriage rate (%). Donut charts below the bubble plots show the proportional composition of seven clonal lineages within each province.

## Discussion

Our comprehensive survey of 6,346 global *K. pneumoniae* isolates reveals that defense systems are not randomly distributed but are structured into lineage-specific profiles that we term "defensotypes." This concept emphasizes the combinatorial architecture and lineage-specific partnerships that define functional immune units [26]. The defensotype framework offers several advantages for understanding *K. pneumoniae* biology. First, it provides a predictive dimension: knowing the ST-KL of an isolate allows inference of its likely defense arsenal, which in turn informs phage susceptibility and plasmid stability predictions [27]. Second, it reveals evolutionary constraints: G1A, G1B, and G1C harbor the highest numbers of defense systems, whereas G1E, G2, and G3 carry fewer, suggesting that high-risk clonal lineages face distinct phage pressures or maintain different plasmid burdens that shape selection for immune investment [28]. Third, it highlights ecological specialization: each major clonal lineage occupies a non-overlapping niche in defense system space, suggesting that successful epidemic lineages have evolved tailored immune strategies rather than converging on a universal optimal repertoire [29].

Co-occurrence network analysis provides strong evidence that defense systems in *K. pneumoniae* operate as cooperative networks rather than independent entities [30]. These systems are organized into lineage-specific functional modules with significant mutual exclusion. This implies that defense systems are not acquired or lost as individual genes but as functionally synergistic units, consistent with the "defense island" concept observed in other bacterial species [31]. The stark contrast in defense system number and composition across lineages suggests widespread evolutionary trade-offs in immune system acquisition. Maintaining multiple active defense modules may impose prohibitive fitness costs [32], and functional redundancy between systems may reduce the marginal benefit of retaining overlapping functions [33]. For instance, G1E-ST23-KL1 entirely lacks AbiE--a system universally present in G1A through G1D--yet compensates with a streamlined DISARM_1-DRT_2-Mokosh_Type_I_A core. Such systematic absence patterns, rather than stochastic gene loss, indicate that defense system composition is shaped by clonal evolutionary history and constrained by functional compatibility and metabolic burden. At the functional level, the retention of toxin-antitoxin systems such as Mok_Hok_Sok and MazEF within specific modules points to abortive infection mechanisms that operate through Type IV toxin-antitoxin modalities [34], operating alongside an expanded arsenal of immune mechanisms [35] that together underscore the remarkable diversity of antiphage strategies in *K. pneumoniae*.

Plasmid association analysis revealed the molecular basis of modular organization and the coupling between defense and resistance mobilomes. Most defense systems were chromosomally localized, reinforcing their role as stable lineage markers. However, notable exceptions illuminate the horizontal dimension: Mok_Hok_Sok was predominantly plasmid-borne in G1E-ST23-KL1 via the IncFIB/IncHI1B cluster (90.6%). The relationship between defense systems and plasmids involves both cooperation and conflict: broad-host-range plasmids carry different defense-resistance payloads across clones, linking phage defense with antibiotic resistance spread, yet these plasmid-borne systems must coexist with chromosomal defenses even though defense systems can sometimes eliminate foreign plasmids [36]. Over time, resistance elements also shape phage-pathogen dynamics [37]. In our data, defense systems rose synchronously with their carrier clones over decades, suggesting that plasmid resistance and chromosomal defense coevolve within expanding lineages, with both contributing to strain fitness. The broad-host-range plasmids may act as flexible vectors that couple phage defense with antimicrobial resistance, potentially through mechanisms involving genetic incompatibility or antagonistic pleiotropy that shape plasmid-host coevolution [38,39].

The tight correlation between defense system prevalence and clonal lineage proportions across time and geography indicates that these signatures can serve as robust epidemiological markers. The accumulation of specific defense systems in high-risk clones supports their utility as readily detectable biomarkers for tracking epidemic lineages [40]. Integrating defense system profiling into existing genomic surveillance frameworks could provide an additional layer of resolution for monitoring clonal emergence and intercontinental spread without requiring exhaustive resistance gene profiling [41]. For instance, the presence of PDC.S15 effectively marks ST11 clones, the absence of AbiE distinguishes the G1E-ST23 hypervirulent lineage, and the RosmerTA-Gabija-CBASS_I module defines the G1C-ST15 emerging threat. Beyond surveillance, the defense system signatures carry direct implications for phage therapy. As phage therapy resurges as an alternative to conventional antibiotics for multidrug-resistant infections, the selection of appropriate phage cocktails requires knowledge of pre-existing defense mechanisms [42]. Clinical implementations have already demonstrated efficacy against recalcitrant *Pseudomonas* infections [43], and our lineage-specific defense atlas provides a predictive foundation for such tailored approaches.

Several limitations warrant consideration. First, our reliance on in silico prediction may miss cryptic or novel systems not captured by current databases [44]. Experimental validation of the co-occurrence modules and their functional synergy remain essential to confirm the protective roles inferred from genomic patterns [45]. Second, geographic sampling biases and the consolidation of rare lineages into broader groups may obscure subtle defense system variations. Third, plasmid association analysis is based on *in silico* detection and may miss plasmid-chromosome interactions or transient plasmid acquisitions. Despite these limitations, the consistency of our findings across population structure, plasmid association, and transmission network analyses supports the robustness of the defense system signatures.

In conclusion, this study establishes that *K. pneumoniae* defense systems are organized into lineage-specific defense system signatures shaped by modular synergy and evolutionary trade-offs. The integration of defense profiles with resistance determinants and plasmid backbones reveals that bacterial immunity is not an isolated trait but an integral component of epidemic clone architecture. As the global threat of multidrug-resistant *K. pneumoniae* intensifies, understanding the immune strategies of successful lineages will be essential for predicting their spread, designing effective phage therapies, and ultimately controlling their clinical impact.

## Methods

### Genome acquisition and quality control

A global genomic dataset of 6,346 *Klebsiella pneumoniae* isolates spanning 60 countries and four decades (1982-2024) was assembled from the PATRIC database. This comprised 1,919 complete genomes and 4,427 whole-genome sequencing (WGS) assemblies. The complete genome set included 1,230 non-Chinese isolates and 689 Chinese isolates; all WGS assemblies were of Chinese origin. Genome quality was assessed using CheckM2 v1.0.0. The following filtering criteria were applied: completeness ≥90%, contamination ≤5%, number of contigs <500, and N50 >10 kb. All 6,346 isolates passed quality control and were used for population structure inference, defense system profiling, and global geographic distribution analyses. The 1,919 complete genomes were used for plasmid association analysis. The 689 Chinese complete genomes, annotated with available metadata including isolation source, collection year, province-level geographic location, and host information (human, animal, environmental, or other), were used for high-resolution transmission network construction and China-specific geographic mapping.

Core genome single nucleotide polymorphisms (SNPs) were identified using Snippy v4.6.0 with *K. pneumoniae* MGH 78578 (GCF_000016305.1) as the reference genome. The core genome alignment was generated from Snippy outputs using SNP-sites v2.5.1. Recombination events were identified and removed using Gubbins v3.0.0 with default parameters. The recombination-filtered core genome alignment was used for all subsequent phylogenetic analyses, including tree reconstruction and pairwise SNP distance calculations. Pairwise SNP distances were calculated from the core genome alignment using custom R scripts.

### Multi-locus sequence typing, capsular typing, resistance and virulence gene identification

All genotypic characterization was performed using Kleborate v2.3.0, a comprehensive tool for *K. pneumoniae* genome analysis. Kleborate was run with default parameters to simultaneously determine: multi-locus sequence type (ST) from the *K. pneumoniae* MLST scheme; capsular (K) and lipopolysaccharide (O) serotypes via integrated Kaptive v2.0.0; antimicrobial resistance (AMR) genes, including carbapenemases, extended-spectrum β-lactamases (ESBLs), and other clinically relevant resistance determinants; and virulence factors, including siderophores (*ybt*, *iuc*, *iro*), hypermucoviscosity determinants (*rmpA*, *rmpA2*), and colibactin (*clb*).

### Population structure and clonal group definition

Population structure was inferred using PopPUNK on all 6,346 genomes in lineage assignment mode (--fit-model lineage, --K 4, --ranks 1,2,3, --threads 32). This generated three hierarchical ranks: rank 1 resolved 146 lineages, with Cluster 1 comprising 71% of isolates. Sub-clustering of Cluster 1 at finer ranks stratified it into five subgroups (G1A–G1E). The remaining isolates were consolidated into G2 (Clusters 2–25) and G3 (Clusters 26–146). Clonal group assignments from PopPUNK were cross-validated with ST-KL typing results from Kleborate.

### Defense system identification and profiling

Bacterial defense systems were identified using a dual-tool approach combining DefenseFinder v1.2.0 and PADLOC v2.0.0. DefenseFinder was employed to detect known defense systems based on curated protein profiles, while PADLOC was used for comprehensive profiling by matching protein sequences against a curated database of >700 Hidden Markov Model (HMM) families of defense system-related proteins. The presence, absence, and subtype classification of all defense systems were compiled into a binary presence-absence matrix for each isolate.

### Defense system conservation and clustering

To evaluate compositional consistency across taxonomic levels, we calculated pairwise Jaccard distances for each analytical unit (clonal group, ST type, and G-ST-KL clonal lineages) based on the binary presence-absence matrix of defense systems. To quantify internal homogeneity, we calculated the coefficient of variation (CV = standard deviation / mean) for each group, where SD is the standard deviation of pairwise Jaccard distances and Mean is the average Jaccard distance within the group.

### Co-occurrence network analysis

Co-occurrence networks were constructed using the Phi (φ) coefficient. For each pair of defense systems, φ was calculated from the presence/absence table. Co-occurrence was considered significant at |φ| > 0.2 with FDR-adjusted *P* < 0.05. Positive φ indicates co-occurrence; negative φ indicates mutual exclusion. Modular structures were identified using the Leiden community detection algorithm (cluster_leiden, igraph R package, resolution = 1.0), with independent module definition for each clonal group.

### Plasmid association analysis

Plasmid sequences were identified using MOB-suite v3.1.0 and PlasmidFinder v2.1, classified by replicon type and incompatibility group. Associations between defense systems and plasmid types were assessed using Fisher’s exact test with FDR correction.

### Phylogenetic signal analysis (Pagel’s λ)

To distinguish vertically inherited from horizontally transferred defense systems, Pagel’s λ was calculated using the phylosig function (phytools R package) for each clonal group with sample size *n* ≥ 5. Pagel’s λ quantifies trait evolution relative to phylogenetic tree topology: λ = 1 indicates perfect phylogenetic signal (vertical inheritance); λ = 0 indicates trait distribution independent of phylogeny (horizontal transfer). Statistical significance was assessed by likelihood ratio tests comparing the fitted λ model against a null model with λ = 0. *P*-values were adjusted using the Benjamini–Hochberg (FDR) method.

### Spatiotemporal and time-series analysis

For global distribution patterns, prevalence (percentage of isolates carrying each system) was calculated for each of the 12 defense systems across 60 countries. Countries with fewer than 10 isolates were excluded. For continental dynamics, isolates were stratified by continent and five-year intervals (2005–2009, 2010–2014, 2015–2019, 2020–2024). Temporal trends were assessed using linear regression (R², p-values). For time-series correlation analysis, annual proportions of each major ST-KL clone were calculated. To enable direct visual comparison, both defense system prevalence and clone proportions were standardized to Z-scores (Z-scores = (x-μ)/ σ), where x is the raw value, μ is the mean, and σ is the standard deviation of the time series. Spearman’s rank correlation coefficient (r) was calculated between each defense system’s annual prevalence and each clone’s annual proportion, with FDR-adjusted p-values.

### Geographic mapping

World maps were generated using the rnaturalearth R package and visualized with ggplot2. For global distribution, pie charts were overlaid on country centroids using the scatterpie R package, with pie size proportional to the square root of isolate count. Countries with fewer than 10 isolates were excluded. Geographic coordinates were projected using the WGS84 coordinate reference system (EPSG:4326).

For the China map, province-level boundaries were obtained from rnaturalearth (*ne_states*). To ensure complete territorial representation, Taiwan provincial boundaries were retrieved separately and merged with mainland China data, with administrative attributes unified to "China". Province names in the metadata were standardized to match map data using a custom mapping dictionary that resolved spelling variants and mapped city-level locations to their corresponding provincial administrative units. Province-level strain counts were mapped to a color gradient (log₁p-transformed) for choropleth visualization. The main panel was projected to 73-136°E and 18-54°N. A South China Sea islands inset (105-123°E, 2-24°N) was generated using *coord_sf* and positioned within the main figure using patchwork (*inset_element*). Defense system carriage rates at each province were visualized as scatterpie charts, with pie radius proportional to the square root of the provincial strain count.

### High-resolution transmission network (China cohort)

Transmission networks were constructed from 689 Chinese complete genomes. Recent transmission was defined as pairwise SNP distance ≤20. Inter-provincial links were defined as isolate pairs from different provinces with SNP distance ≤20. For each province pair, pair_count (number of connecting isolate pairs) and avg_SNP (mean SNP distance) were calculated. Networks were visualized as weighted undirected graphs (igraph R package), with nodes representing provinces, edges representing links, edge width proportional to pair_count, and node size proportional to isolate count. Node degree (connected provinces) was calculated to identify network hubs.

## Statistical analysis

All analyses were performed using R v4.3.0. Continuous variables were compared using Kruskal-Walli’s test (multi-group). Categorical variables were compared using Fisher’s exact test or chi-square test. All p-values were two-sided and adjusted for multiple testing using Benjamini-Hochberg (FDR) correction. P < 0.05 was considered statistically significant.

## Data availability

All genome sequences used in this study are publicly available from PATRIC. Accession numbers and metadata are provided in Supplementary Table S1.

## Author contributions

**Xiaofu Wan:** Conceptualization, Investigation, Visualization, Formal analysis, Writing-original draft, Writing-review & editing. **Liyin Ji & Shuhong Han & Zixun Lin**: Investigation. **Yukun Zeng & Min Ming & Wenheng Gan**: Formal analysis. **Xiangke Duan**: Resources, Supervision. **Hongzhou Lu:** Funding acquisition, Resources, Supervision. **Jiayin Shen**: Conceptualization, Investigation, Writing -Original draft preparation, Writing - Review & Editing, Project administration, Funding acquisition. All authors discussed the results, commented on and approved the final manuscript.

## Acknowledgments

This study was supported by National Natural Science Foundation of China (No. 82473753,No. 82070420); National Key R&D Program of China (No. 2025YFC3408400); Prevention and Control of Emerging and Major Infectious Diseases-National Science and Technology Major Project (No. 2025ZD01908400); Shenzhen key laboratory for infectious diseases; Shenzhen Clinical Medical Center for Emerging infectious diseases (LCYSSQ20220823091203007); Shenzhen Science and Technology Program (No. KJZD20240903100500001), Shenzhen Natural Science Foundation (No.JCYJ20250604143818024) and Shenzhen High-level Hospital Construction Fund.

## Competing interests

The authors declare no competing financial interests.

## Notes

### Competing Interest Statement

The authors have declared no competing interest.

### Summary of Updates

This version of the manuscript has been revised to update the following: (1) Corrected the administrative boundary matching in the geographic visualization (Figure 6 ) to ensure standardized representation of national territory. (2) Optimized the inset map for the South China Sea islands by enlarging the display area, increasing line clarity, and adding clearer border annotations to improve resolution and visual recognition. (3) Minor text edits in the Methods section to clarify the spatial data processing workflow. (4) Supplemental files updated.

