## Supplementary Figures for "Exploratory profiling of defense system signatures in *Klebsiella pneumoniae* clonal populations"

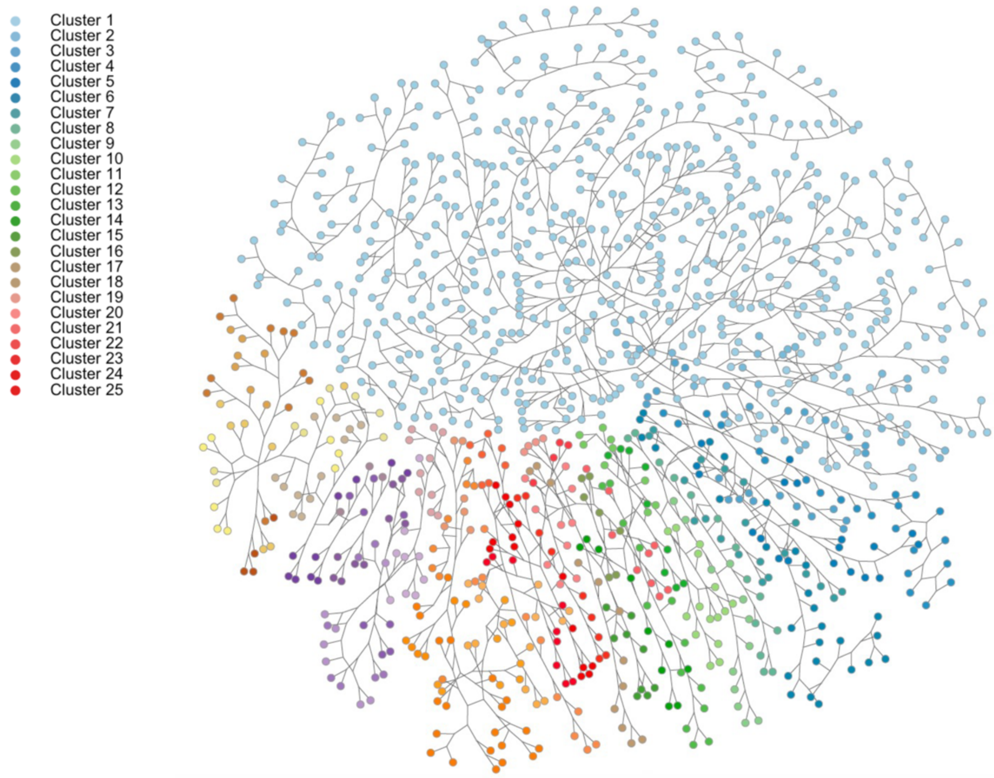


**Figure S1. PopPUNK clustering of a random subset of 1,100 *K. pneumoniaegenomes.***

A random subset of 1,100 genomes from Clusters 1-25 was selected for PopPUNK clustering analysis. Each node represents a genome, with colors denoting cluster assignments.

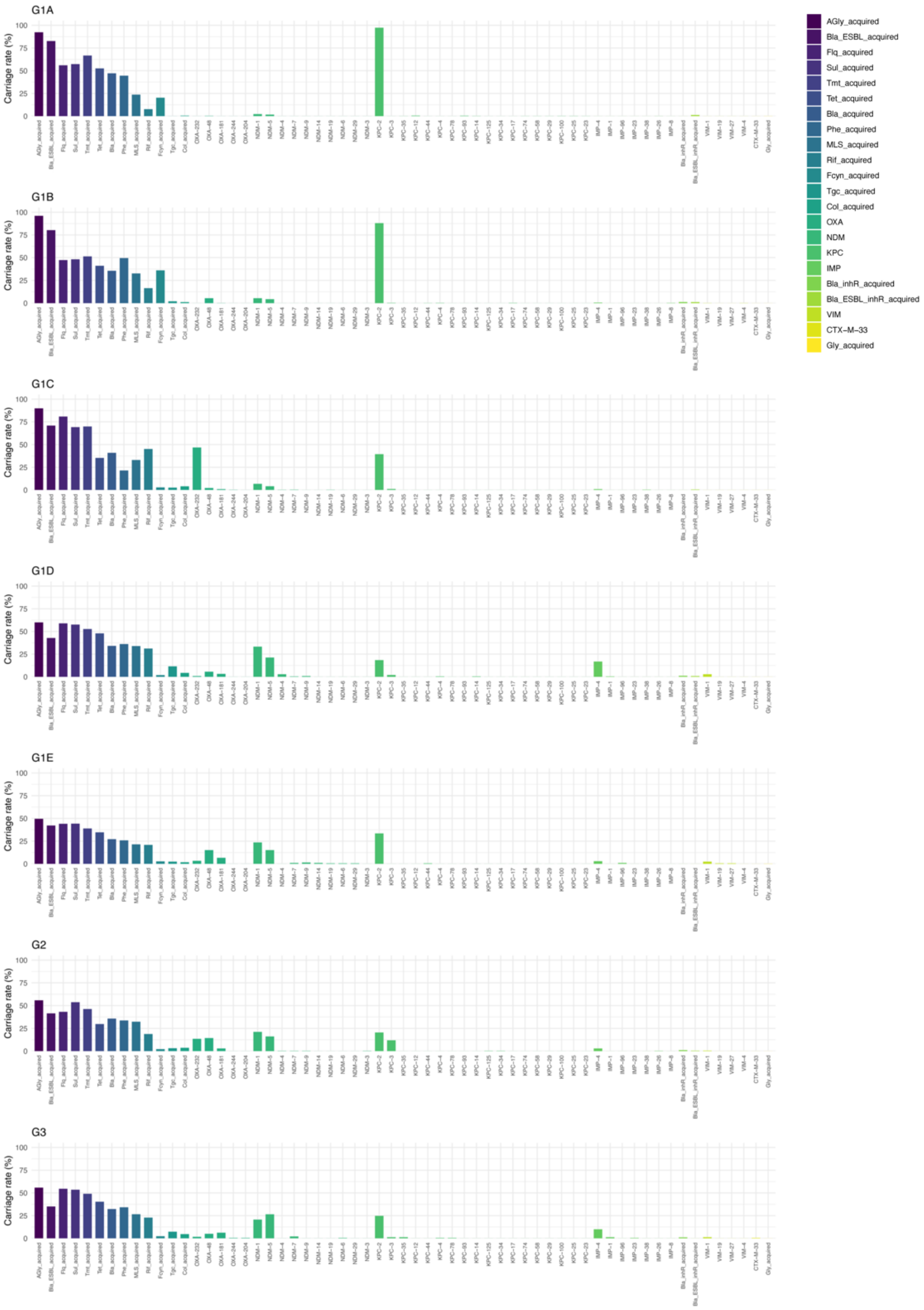


**Figure S2. Resistance gene carriage rates across the seven groups.**

Bar plots showing the carriage rate of individual resistance genes within each of the seven groups.

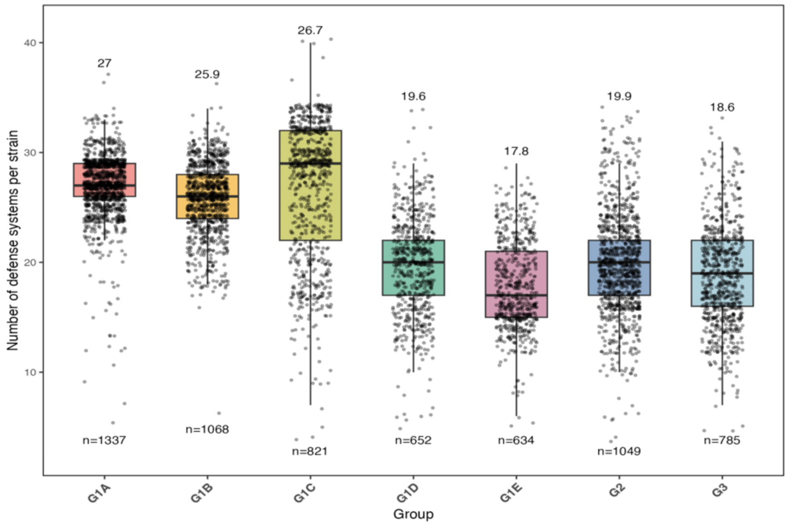


**Figure S3. Defense system counts across seven groups.**

Box plot showing the distribution of the number of defense systems per strain across the seven groups (G1A-G3).

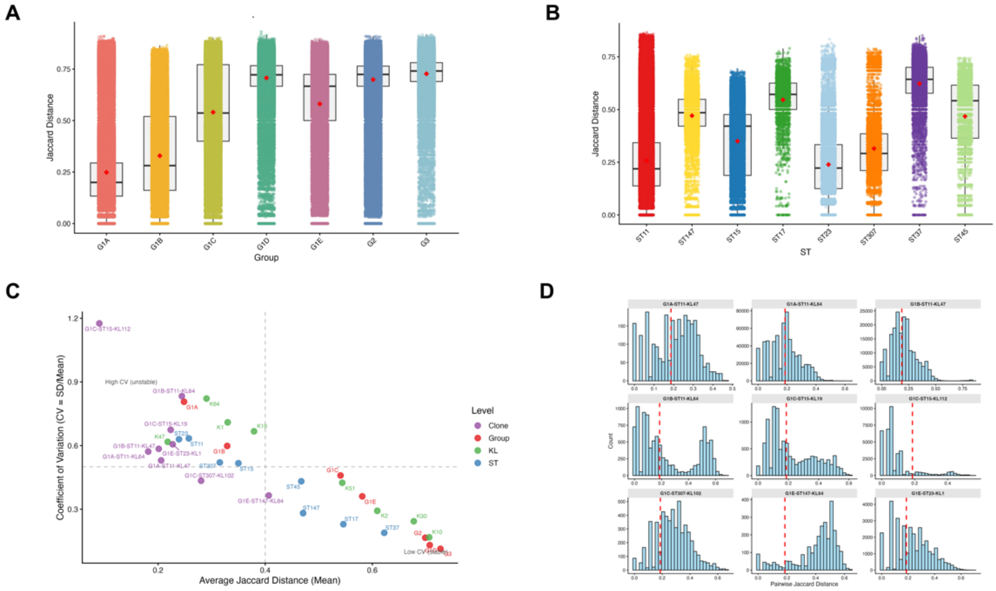


**Figure S4. Compositional conservation of defense systems across hierarchical levels.**

**(A)** The distribution of pairwise Jaccard distances within each of the seven groups.

**(B)** The distribution of pairwise Jaccard distances within the eight most prevalent ST types (ST11, ST15, ST23, ST147, ST37, ST307, ST17, and ST45).

**(C)** Scatter plot of average Jaccard distance (Mean) versus coefficient of variation (CV = SD/Mean) across groups at the Group, ST, KL, and clonal lineage levels. Each point represents a lineage, with point color indicating the hierarchical level. Lines represent CV = 0.5 and Mean = 0.4 thresholds.

Lineages in the bottom-left quadrant (low Mean, low CV) represent the most stable and conserved defense system signatures.

**(D)** Histogram of pairwise Jaccard distances within the clonal lineage.

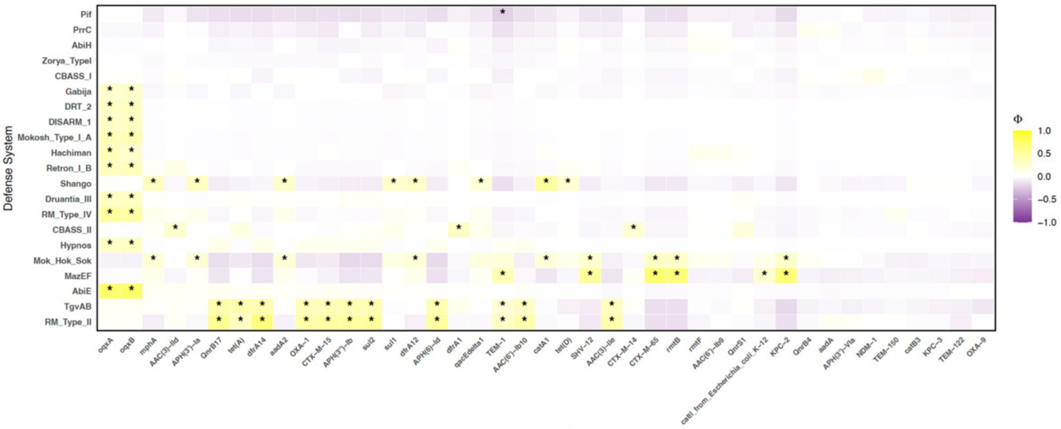


**Figure S5. Plasmid-mediated associations between defense systems and resistance genes.** Co-carriage heatmap showing the prevalence of defense system-resistance gene pairs on plasmid clusters across the complete gene of *K. pneumoniae*. Cells are color-coded from purple (negative association) to yellow (positive association), with color intensity proportional to Phi. *Asterisks indicate the significance level after FDR correction: *Q < 0.05.

composition of seven clonal lineages within each province.

.
